# Mapping local and global liquid-liquid phase behavior in living cells using light-activated multivalent seeds

**DOI:** 10.1101/283655

**Authors:** Dan Bracha, Mackenzie T. Walls, Ming-Tzo Wei, Lian Zhu, Martin Kurian, Jared E. Toettcher, Clifford P. Brangwynne

## Abstract

Recent studies show that liquid-liquid phase separation plays a key role in the assembly of diverse intracellular structures. However, the biophysical principles by which phase separation can be precisely localized within subregions of the cell are still largely unclear, particularly for low-abundance proteins. Here we introduce a biomimetic optogenetic system, “Corelets”, and utilize its rapid and quantitative tunability to map the first full intracellular phase diagrams, which dictate whether phase separation occurs, and if so by nucleation and growth or spinodal decomposition. Surprisingly, both experiments and simulations show that while intracellular concentrations may be insufficient for global phase separation, sequestering protein ligands to slowly diffusing nucleation centers can move the cell into a different region of the phase diagram, resulting in localized phase separation. This diffusive capture mechanism liberates the cell from the constraints of global protein abundance and is likely exploited to pattern condensates associated with diverse biological processes.

## Introduction

In addition to the canonical vesicle-like membrane-bound organelles, there are dozens of different types of organelles that are not membrane-bound – from the nucleolus and stress granules to processing bodies and signaling clusters. These represent dynamic molecular assemblies, which can play numerous roles in sequestering biomolecules, facilitating reactions, and channeling intracellular signaling.

Recent studies on membrane-less organelles have revealed that their assembly arises from liquid-liquid phase separation, driven by weak multivalent interactions often involving intrinsically disordered protein regions (IDPs/IDRs) and nucleic acids (Brangwynne et al. 2009; Li et al. 2012; Nott et al. 2015; Brangwynne et al. 2011; Elbaum-Garfinkle et al. 2015). These interactions give rise to intracellular condensates, which represent stable coherent organelles that nonetheless typically exhibit dynamic molecular exchange and liquid phase fluidity (Shin & Brangwynne 2017; Banani et al. 2017).

In many cases these membrane-less condensates are spatially patterned within living cells. For example, germline P granules are known to form via a liquid-liquid phase-separation process that is modulated across the anterior-posterior embryo axis, giving rise to an asymmetric localization implicated in early cell fate specification (Brangwynne et al. 2009; Smith et al. 2016). The nucleolus is a multi-phase nuclear body that assembles at specific genomic loci associated with ribosomal RNA transcription (Feric et al. 2016; Berry et al. 2015). The nucleolus is increasingly viewed as a hypertrophied example of phase-separated foci that form at particular genomic loci (Zhu & Brangwynne 2015), including other transcriptionally-active genes (Hnisz et al. 2017), or at regions of transcriptionally-inactive heterochromatin (Larson et al. 2017; Strom et al. 2017). RNA accumulation (Berry et al. 2015), chemical reactivity (Zwicker et al. 2014), or morphogen gradients (Brangwynne et al. 2009), have been proposed to drive patterned phase separation. Nevertheless, it is still unclear how IDPs and other sticky ligands distributed throughout the cell, often at relatively dilute concentrations, can be rapidly and controllably induced to condense at particular subcellular locations in the cell.

Elucidating intracellular phase behavior has been challenging due to the lack of tools for triggering, shaping, or destabilizing condensates in their native cellular environment. Recently, we developed a photo-activated system for reversibly controlling IDR-driven phase transitions using photo-oligomerizable Cry2 proteins (Shin et al. 2017). This system showed a threshold saturation concentration for phase separation, and exhibited other behaviors consistent with a classic liquid-liquid phase separation, which may be linked to subsequent gelation transition. However, the uncontrollable oligomerization of the Cry2 clusters did not allow for quantitative interrogation of phase behavior; furthermore, the slow deactivation timescale (∼10 minutes) fails to facilitate tight spatial control of phase separation. Another recent study focused on light and chemically-activated multimerized interaction domains, which gave rise to intracellular gel-like structures (Nakamura et al. 2017). But to date, no tools have enabled mapping intracellular phase diagrams, hindering our understanding of the local and global phase behavior within living cells.

To address this gap, we developed an optogenetic system inspired by endogenous molecular architectures in which oligomerization domains often recruit IDR-rich proteins. For example, nascent ribosomal RNA (rRNA) transcripts (Falahati et al. 2016; Berry et al. 2015) or long non-coding RNA (lncRNA) such as Neat1 (West et al. 2014), and other types of RNA are associated with specific DNA loci may serve as scaffolds for locally enriching self-interacting IDPs (Dundr & Misteli 2010); DNA itself could also serve as an oligomerization platform for promoting local transcriptional condensates (Hnisz et al. 2017). Finally, a number of proteins play scaffolding roles, for example the nucleolar protein Npm1, which pentamerizes to form a radial array of IDRs and RNA-binding domains necessary to promote phase separation at the heart of nucleolar assembly (Feric et al. 2016; Mitrea et al. 2016). Inspired by these native molecular architectures, we reasoned that an approach to precisely control the oligomerization state of IDPs could elucidate the underlying biophysical mechanisms by which intracellular phase transitions are locally controlled in cells.

## Results

### Corelets enable light-activated intracellular liquid droplet condensation

To parse the effect of multivalent scaffolding of IDRs on intracellular phase separation, we developed Corelets, an optogenetic system which mimics the radial architecture of Npm1 pentamers, using a light-activatable high valency core. The core is comprised of 24 human ferritin heavy chain (FTH1) protein subunits, which self-assemble to form a spherical particle of 12 nm diameter (Bellapadrona & Elbaum 2014). We fused FTH1 to a nuclear localization signal (NLS) and an engineered protein iLID, which strongly heterodimerizes (Kd ∼ 130 nM) with its cognate partner, SspB, in response to blue light activation (Guntas et al. 2015) (Figure 1A). SspB was fused to various IDRs, as well as full length IDR-containing proteins, implicated as drivers of intracellular phase separation, such that the ferritin core would serve as a well-defined multivalent scaffold for light-activated IDR oligomerization.

**Figure 1.**
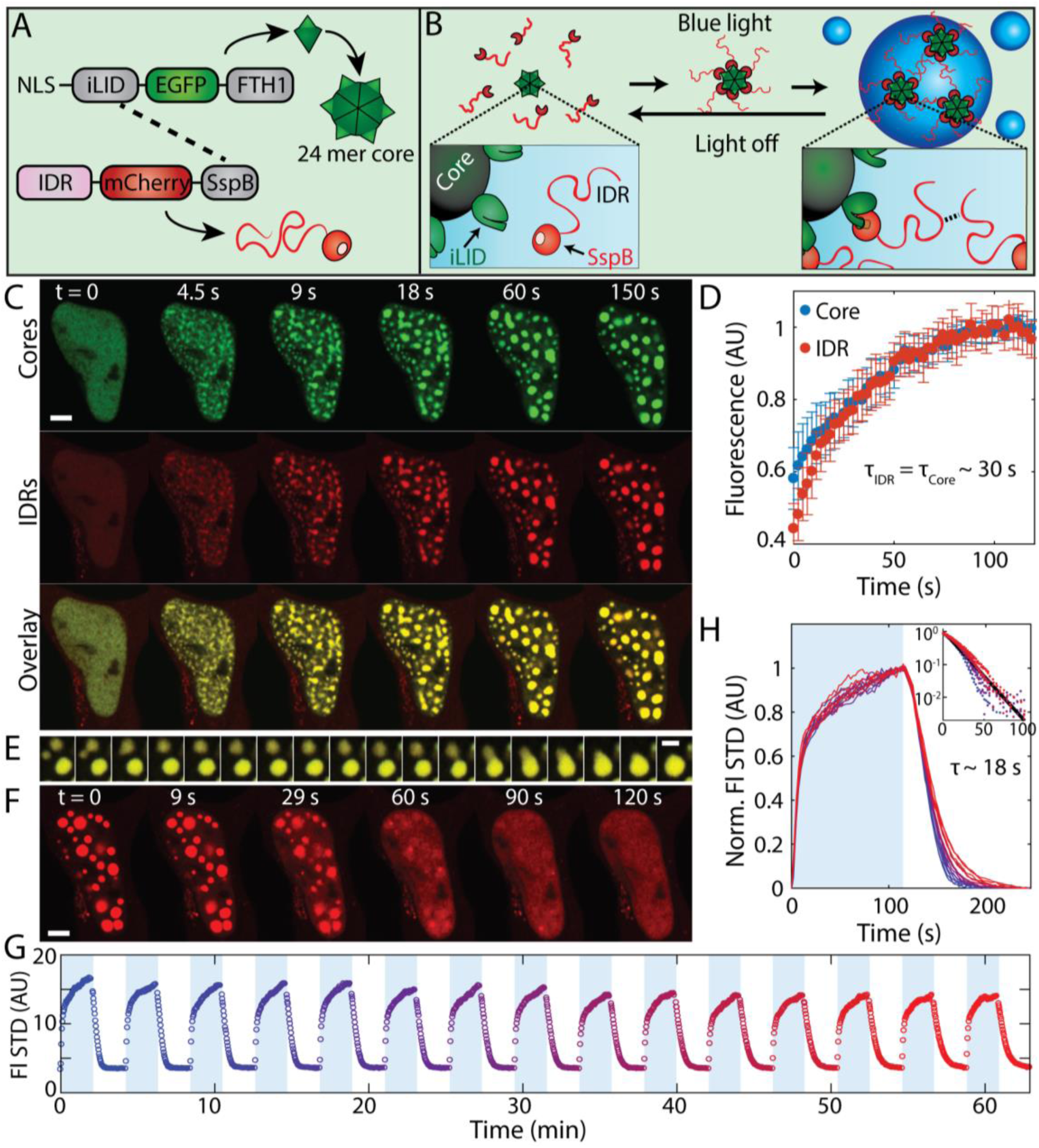
Corelets enable light-activated liquid droplet condensation. **(A)** Schematic diagram of Corelet system. Corelets consists of two modules: first, a nuclear targeted GFP-ferritin core functionalized by 24 photo-activatable iLID domains and second, iLID’s cognate partner, SspB, mCherry-labelled and conjugated to an IDR, such as FUS_N_. **(B)** Schematic of Corelet phase separation. Upon blue-light activation, up to 24 IDR domains are captured by Ferritin cores, which subsequently phase separate in a reversible manner. **(C)** Time lapse confocal imaging of photo-activated Corelet-expressing HEK293 cells. Images show phase separation and colocalization of cores (green) and IDRs (red). **(D)** Corelet condensates exhibit liquid-like properties as inferred by rapid fluorescence recovery after photo-bleaching of both cores (red) and IDR (blue) components (error bars are standard deviation of the mean), and **(E)** rapid fusion and coarsening (4.4 s between frames). **(F)** After 10 min of activation, condensates disassemble within ∼0.5-2 min of blue-light being turned off. **(G)** Change in standard deviation of IDR fluorescence, which reflects transitions in spatial heterogeneity of composition,, indicates full reversibility during 15 on-and off cycles (4 min each). **(H)** Overlay of data from G, showing little change in condensation and de-condensation dynamics over multiple cycles. Inset, overlaid disassembly dynamics on semi-log plot, indicating a dissociation rate of ∼18 s. Bars are 5 µm for C and F and 2 µm for E.

In response to blue light activation, as many as 24 IDRs are induced to directly decorate the Ferritin core, thus forming rapidly responsive self-interacting protein assemblies. We first utilized an N-terminal FUS IDR (FUS_N_) fused to SspB (Figures 1A and 1B). For this FUS_N_ Corelet system, condensation is apparent within ∼1-2 seconds after blue light illumination and saturates within a few minutes (Figures 1C and S1; Movie S1). These FUS_N_ Corelet condensates are liquids, as apparent from their rapid and complete fluorescence recovery after photobleaching (FRAP) (Figure 1D), and their ability to rapidly fuse with one another and round up upon contact (Figure 1E). When activating illumination is turned off, the droplets quickly dissolve back to a uniform phase (Figure 1F). Moreover, when we applied sequences of uniform blue light activation cycles, FUS_N_-Corelets could be repetitively assembled through dozens of on-off cycles, with little apparent change in the disassembly kinetics (Figures 1G and 1H). A unique feature of Corelets is that while the strong light-driven interactions effectively switch the system into a one component system of IDR coated cores, they do not directly contribute to the cohesive interactions of the emergent liquid phase, which instead rely exclusively on homotypic IDR-IDR interactions. Consistent with IDR-driven Corelet phase separation, we observe no phase separation in control constructs that do not have IDRs (Figure S1), and we find significant recruitment of SspB-free full-length FUS and FUS_N_ proteins to FUS_N_-Corelets (Figure S2).

### Corelets drive phase separation with multiple IDRs and in multiple living systems

Liquid corelet condensates could be formed not only in the nucleus, but also in the cytoplasm, by excluding the NLS from Ferritin constructs (Figure 2A). Full length FUS can also be utilized (Figure 2B), which will have likely applications in generating synthetic versions of endogenous condensates such as stress granules or transcriptional foci. Moreover, similar phase separated liquid condensates could also be formed from Corelets comprising a number of different IDR-containing constructs. These include IDRs from other RNA binding proteins associated with stress granules, such as hnRNPA1_C_ and TDP43_C_, as well as the germ granule components DDX4_N_ and PGL1 (Figures 2C-2F). Interestingly, the nature of the condensed phase varies with the type of IDR used; while FUS_N_, hnRNPA1_C_, and DDX4_N_ Corelets display spherical shapes, full-length FUS and TDP43_C_ Corelets display more irregular, slowly coarsening structures, apparently reflecting a more solid-like character due to stronger homotypic interactions. Corelets can be dynamically assembled not only in various cultured cell lines including HEK293, NIH3T3, and U2OS (Figures 1C, S1, and 2C, respectively) without observable interference by endogenous Ferritin (Figure S3), but also in *Saccharomyces cerevisiae* (Figure 2F) and *Caenorhabditis elegans* (Figure 2G).

**Figure 2.**
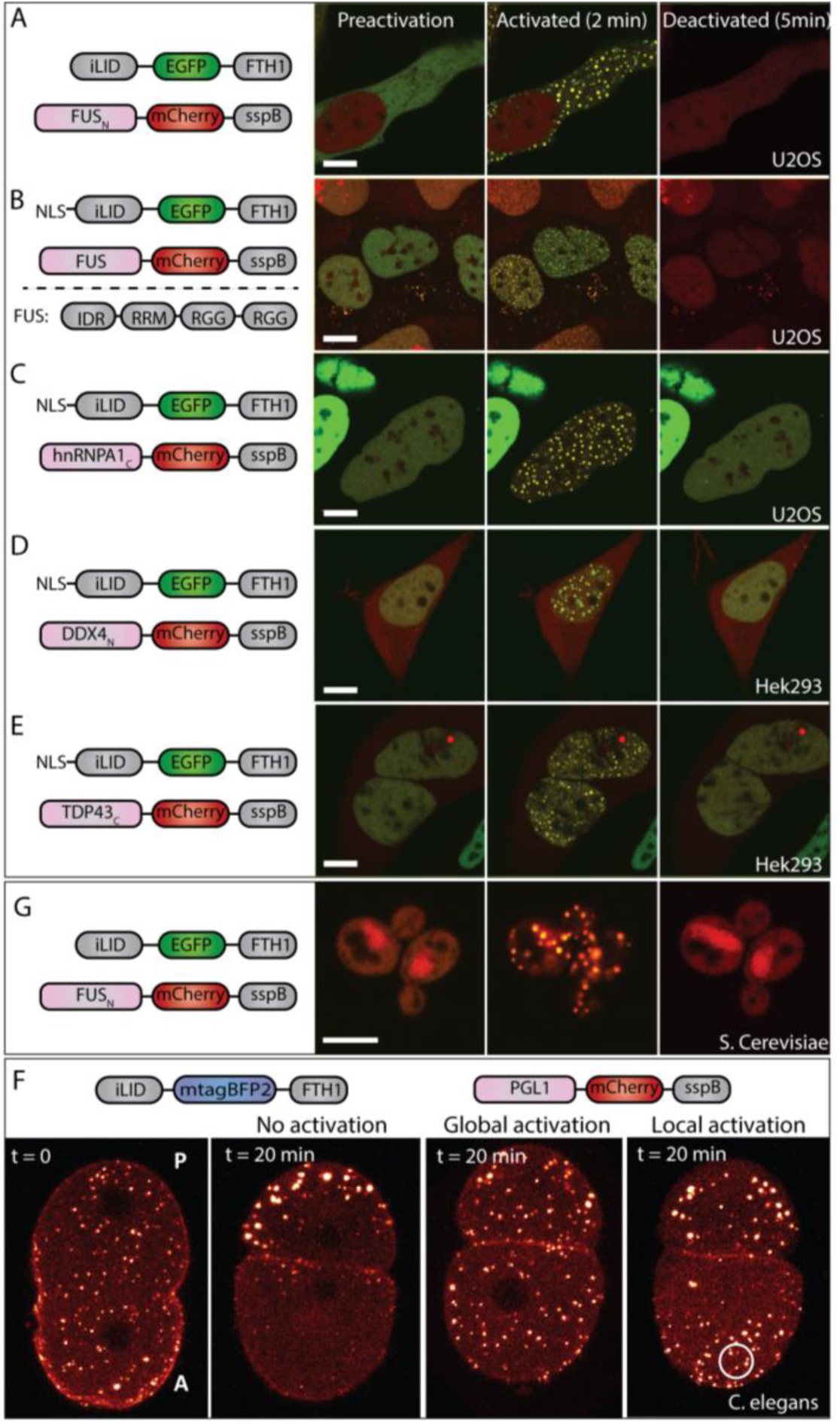
Corelets drive phase separation with various IDRs and in various living systems. Fluorescence images of representative stable cells expressing nuclear Corelets with NLS free cytoplasmic Corelets with **(A**,**F)** FUS_N_ IDR, **(B)** full length FUS protein, and **(C)** hnRNPA1_C_, **(D)** DDX4_N_, **(E)** TDP43_C_ IDRs. A-B are U2OS cells, C-E are HEK293, and F is *S. cerevisiae*. Images taken before and after 2 min of activation and 5 min after deactivation. All images show overlay of GFP (core) and mCherry (IDR/IDP) except light off (t=7min) images for A, B, and F, which show mCherry signal only. All constructs display reversible puncta formation with various disassembly times. Condensates formed by full length FUS and TDP43_N_ Corelets display slower coarsening and appear to be less spherical compared to condensates made by FUS_N_, hnRNPA1_C_, and DDX4_N_ Corelets. We note that cytoplasmic FUS_N_ Corelets display higher mobility as compared to nuclear FUS_N_ Corelets. Schematic diagram of FUS domains are shown in B. **(G)** Local and global activation of PGL-1 Corelets in *C. elegans* one-cell embryo. Images shown as heat colormap of mCherry signal. Prior to pro-nuclei meeting (left), PGL1-SspB components are recruited by the native P granules, which are distributed uniformly throughout the embryo at this early time point. Without any light activation, P granules segregate to the posterior end (P). Under global activation, Corelet puncta appear throughout the embryo. When local activation is applied, native granules persist on the posterior end, while PGL1-Corelets puncta emerge nearby the activated region (solid circle). Schematic diagrams of Corelet constructs are shown. Scale bars is 5 µm.

### Full Phase Diagram of Globally Activated FUS_N_ Corelets

To quantitatively test whether the observed transition indeed corresponds to liquid-liquid phase separation, we analyzed cells with different relative expression levels of the FUS_N_-SspB (“IDR”) and 24-mer Ferritin-iLID (“Core”) components; we define their average nuclear concentration ratio as 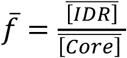. Calibrated pixel intensity histograms show a unimodal distribution of the core concentration before activation, while after activation phase separating cells exhibit broadened and even bimodal distributions, with the two peaks becoming farther apart for cells with high 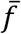 (Figures 2A-C). For these global activation experiments, the molar ratio inside droplets *f*_*Dense*_ and outside droplets *f*_*Dilute*_ are very close, and similar to 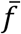 (Figure 3D), consistent with dynamic exchange of IDR-bound Ferritin cores and very low concentrations of unbound FUS_N_–SspB (Figure S4). However, as 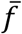 approaches the binding capacity of cores, this correspondence begins deviating, suggesting the buildup of an unbound IDR population that partitions asymmetrically.

**Figure 3.**
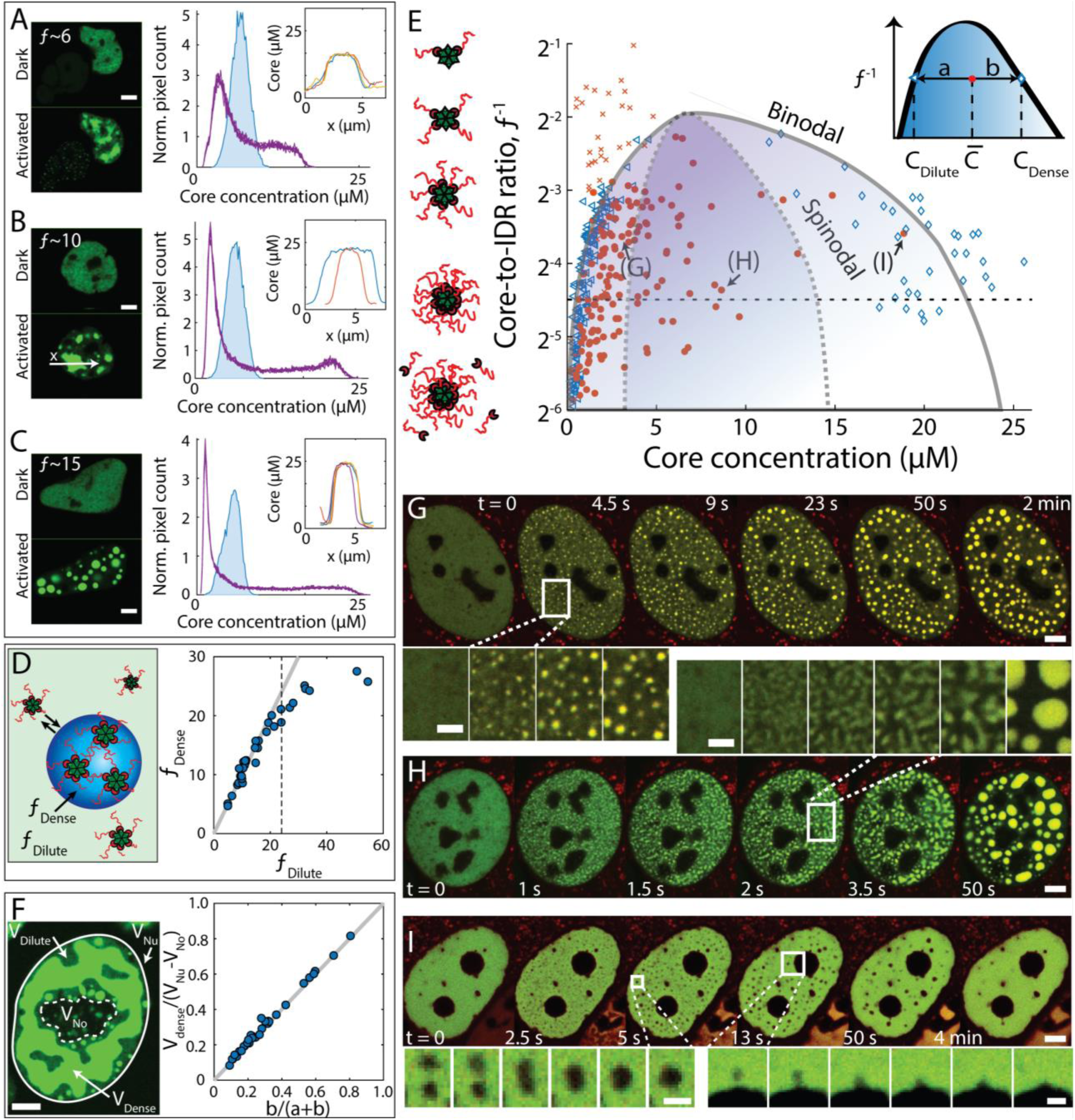
FUS_N_ Corelets follow liquid-liquid phase separation phenomena. **(A-C)** Fluorescence images and histograms of Core concentration before (blue) and after (purple) 10-minute photoactivation, for three representative cells with similar 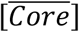, but different 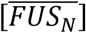, and thus also a different ratio 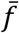. Cells with larger 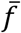 yield a greater separation between concentration inside droplets, [*Core*]_*Dense*_, and outside droplets, [*Core*]_*Dilute*_. Insets show profiles of a small number of droplets. Large droplets in every cell reach similar plateau values. **(D)** Despite major concentration differences, concentration ratios in dilute *f*_*Dilute*_ and dense *f*_*Dense*_ phases are similar (compared to gray line of slope 1) as long as cores are not saturated (dotted line), suggesting that within this range the activated Corelets behave as a single component phase-separated system. **(E)** Phase diagram of FUS_N_ Corelets, where solid red symbols indicate average nuclear concentrations for which phase separation is observed, and ‘x’ red symbols are concentrations where no phase separation is observed. Blue triangles and diamonds indicate the valency/concentration of dilute phase and dense phase, respectively. Dashed horizontal line corresponds to fully-coated Corelets with 24 IDRs per Ferritin Core. Vertical axis on a log 2 base. Binodal and spinodal lines were determined accordingly with the observed mode of transition as shown in Figure S5. **(F)** Volume fractions predicted by lever-rule (see inset of panel E for definition of “a” and “b”) are consistent with volume fraction segmentation of dense (V_Dense_, bright green) and dilute phases (V_Nu_-V_No_-V_dense_, dark green), where V_Nu_, V_No_, and V_Dense_ represent the relative confocal volume of the nucleus (full line), nucleolus (dashed line), and dense phase (bright green) respectively. **(G)** Nucleation and growth of Corelets observed at concentrations near the binodal line. **(H)** Spinodal decomposition observed deep within the binodal. Insets for G-H show the differing morphologies. **(I)** Nucleation and growth of a dilute phase within dense FUS_N_ Corelets phase was observed at high concentration. Insets for I show time lapse images of fusion between nucleated dilute droplets (left) and between dilute droplet and a nucleolus (right); frame interval=0.5s. Bars are 2 µm for enlarged images in G-I and 5 µm elsewhere.

For cells with very low 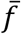, phase separation never occurs, while cells with higher 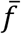 typically do form condensates (Figure 3E, x and o symbols respectively), consistent with 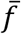 representing an effective interaction energy between IDR-decorated cores. These data reflect the position of the cell with respect to a concave-down binodal phase boundary, as seen by plotting the inverse molar ratio 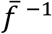, against 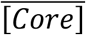 (Figure 3E). As expected for a binodal phase boundary, the concentration of Cores measured outside of the droplets, [*Core*]_*Dilute*_, agrees well with the left-arm of the boundary between non-droplet forming cells and droplet-forming cells (Figure 3E). Moreover, the right arm of the binodal can be determined from the protein concentration in droplets (Wei et al. 2017), in this case [*Core*]_*Dense*_, which at high 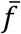 corresponds to a mean spacing between Cores of roughly 40 nm; we estimate that Corelet components occupy ∼10% of the condensate volume. Our determination of the location of the binodal is further supported by the lever rule for the volume fraction of droplets (Figure 3F). Consistent with phase separation theory (Berry et al. 2018), we find that cells expressing concentrations deep within the two-phase region (i.e. highly supersaturated) exhibit early stage coarsening morphologies associated with spinodal decomposition, while more moderately supersaturated cells exhibit nucleation and growth (Figures 3G and 3H; Movies S2A and S2B). Moreover, for some highly expressing cells, we observe spinodal decomposition or nucleation and growth of dilute-phase droplets within a continuous condensed phase, as expected on the right half of the two-phase regime of the phase diagram (Figure 3I; Movies S2C and S2D). These dilute phases fuse to one another, coarsen, and further fuse to nucleoli and nuclear lamina, while yielding condensates that may occupy a volume of over 70% of the nuclear volume. Taken together, the various condensation modes allow us to determine the approximate location of the spinodal boundary (Figures 3E and S5).

### Local concentration amplification of IDRs occurs through diffusive IDR capture by slowly diffusing cores

In the experiments described above, we define the activation zone over the entire nucleus of the cell under study. However, occasionally only a fraction of adjacent cell nuclei were included in the activation zone. We noticed that in half-activated nuclei, droplets appear to be significantly larger close to the border between illuminated and non-illuminated fractions of the nucleus (Figure 4A). In cells illuminated with low power light, this effect becomes less prevalent, and the droplet size and number tend to be more evenly distributed throughout the activation zone, as shown for the half-illuminated cells expressing hnRNPA1_C_ Corelets in Figure 4B. However, when we brightly half-illuminate nuclei with low 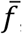, we find that droplets form in a tight line at the illumination boundary (Figure 4B). Similar findings are observed with FUS_N_ (Figure 4C). Prior to droplet nucleation, the IDR concentration exhibits a peak at the boundary, with depletion into the non-illuminated region (Figure 4D).

**Figure 4.**
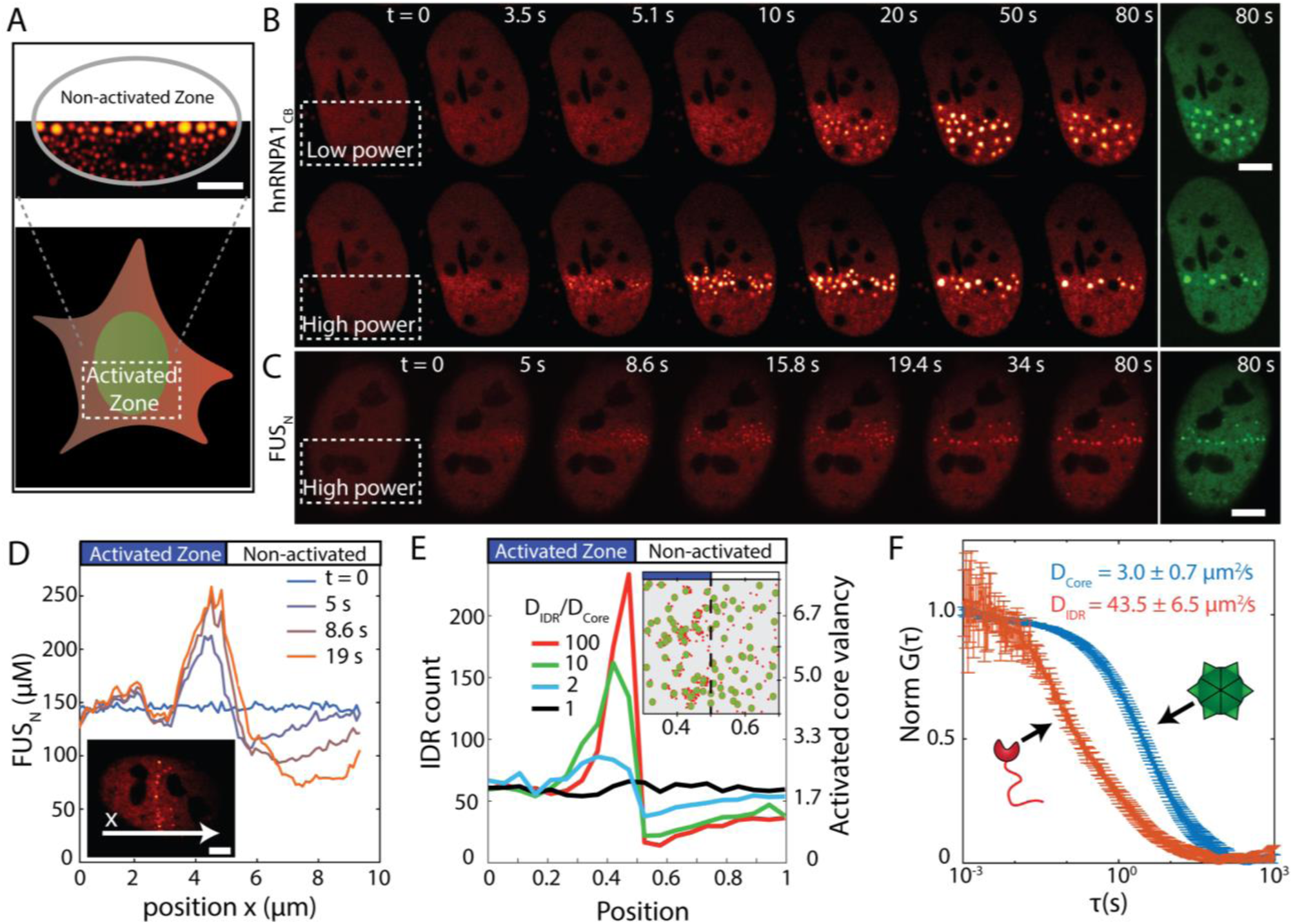
Concentration amplification of IDRs occurs through diffusive capture by slowly diffusing cores. **(A)** Activating a fraction of a cell nucleus leads to non-uniform droplet size-distribution. **(B)** Time-lapse imaging of cell nuclei partially activated by low (top) or high (bottom) activation power. Low power yields uniform nucleation within activated zone, while high power yields preferential nucleation at the boundary between activated and non-activated zones for both hnRNPA1_N_ and **(C)** FUS_N_ based Corelets. Red panels are IDR channel, and green panels are Core channel. **(D)** Time-dependent concentration profiles across cell nucleus with the onset of high power photo-activation. FUS_N_ molecules progressively accumulate at the activated zone boundary, and are depleted within the non-activated zone. **(E)** Simulation demonstrating that capture of IDRs by multivalent cores are sufficient for local enrichment at the activation interface, but only if the Cores diffuse more slowly than IDRs. Inset showing snapshot of a simulation with *D*_*IDR*_/*D*_*core*_ ≅ 10. **(F)** FCS normalized autocorrelation plots measured for core (blue) and IDR (FUS_N_, red) components in the nucleoplasm. The measured diffusion coefficients, *D*_core_ = 3.0 ± 0.7 *μm*^2^/*s* and *D*_*IDR*_ = 43.5 ± 6.5 *μm*^2^/s, were determined by fitting with a simple translation diffusion model. Bars are 5 µm.

We reasoned that the IDR buildup at the illumination interface could occur because the cores at the interface are accessible to and can readily capture IDRs diffusing in from the non-illuminated region. To quantitatively examine this physical picture, we developed a simple computational simulation in which IDRs are modeled as particles that can adhere to the surface of a larger core particle, which supports up to 24 bound IDRs (SI). For low 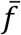, activation of only half of the cell results in a large buildup of IDR particles at the activation interface, with a depletion in the non-activated region, exactly as in experiments (Figure 4E; Movie S3A). Interestingly, the simulation suggests that this effect depends on the relative diffusivities, *D*, of the core and IDR particles: for higher ratios of *D*_IDR_/ *D*_Core_, we find a significant buildup, while for *D*_IDR_=*D*_Core_ we find no buildup (Figure 4E). Using fluorescence correlation spectroscopy (FCS) to measure the Ferritin and IDR diffusivities in living cell nuclei, we find that *D*_FUS_ =43.5+/-6.5 µm^2^/sec, while the core diffusivity is significantly less, even without bound IDRs: *D*_Core_ =3.0+/-0.7 µm^2^/sec, such that *D*_IDR_/ *D*_Core_> 10 (Figure 4F). These data suggest that Ferritin cores act not only as multimerizing cores, but upon local activation can serve as slowly diffusing IDR sinks. These cores thereby capture the pool of IDRs diffusing in from the non-illuminated side of the nucleus, slowing their diffusion, and locally enriching them to drive phase separation. The more uniform IDR buildup and droplet condensation under weaker illumination (Figure 4B) thus results from the associated lower binding capacity of Ferritin cores, which therefore saturate at lower valency and allow IDRs to propagate deeper into the activated region, consistent with simulations in which cores can only bind a small number of IDRs (Movie S3B).

### Phase separation in highly dilute cells upon local scaffold activation

Since partial activation of the cell can give rise to gradients in IDR concentration and valency, we wondered what would happen with non-phase separating cells. In order to be sensitive to differences in local concentration, we first chose a cell with Core and IDR concentrations that position it close to the upper critical point on the phase diagram, such that spatial concentration fluctuations, but not distinct condensates, are observed upon uniform illumination (Figure 5A; Movie S4). Consistent with diffusive capture and local concentration amplification, under half-cell activation, the nucleus indeed exhibits small distinct condensates (white arrows, Figure 5A). Moreover, for 1/3, 1/4, and 1/6 nuclear area activation, larger droplets are observed to condense in the activated region (Figure 5A; Movie S4). Interestingly, as a smaller fraction of the nucleus is illuminated, the molar ratio inside droplets, *f*_*Dense*_ is no longer the same as the average in the entire nucleus, 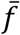. Together with measurements of [*Core*]_*Dense*_, we find that the droplets now correspond to points still on the binodal curve, but much deeper within the two-phase region (Figure 5B). Moreover, regions outside of the activation zone now correspond to much lower values of 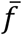 (Figure 5B). Thus, activating local regions of the cell gives rise to diffusive IDR capture and amplification of valence/concentration, causing local supersaturation and droplet condensation, even under globally dilute IDR concentrations.

**Figure 5.**
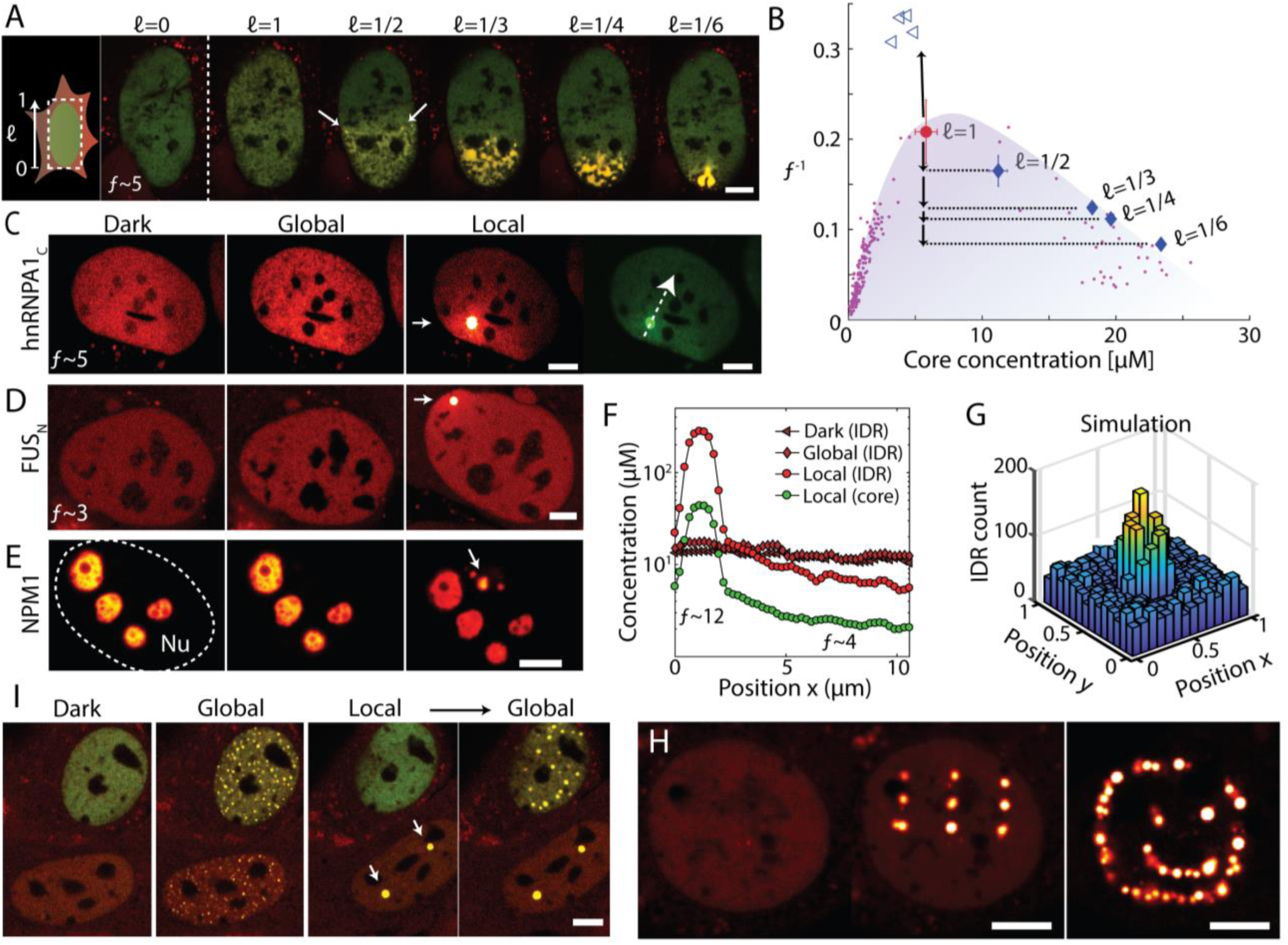
Local but not global phase separation by diffusive capture. **(A)** Performing multiple on-off cycles on subfractions of a near-critical cell expressing FUS_N_ Corelets gives rise to a gradual transition from fluctuating coexisting phases under full nucleus activation (*ℓ*=1) into clear phase separation with increasingly concentrated condensates as size of activated zone decreases. **(B)** The smaller the activated zone, the deeper the cell locally plunges into the two-phase region. When mapped according to local valency and core concentration, resulting condensates that follow the binodal phase boundary. Purple points represent the binodal-line forming data points shown in Figure 3E. Open triangles correspond to non-activated regions of the nucleus. Note that vertical axis is in linear scale. **(C-E)** Photo-activating a 0.5 µm spot in globally non-activatable cells, expressing either **(C)** hnRNPA1_N_ Corelets, **(D)** FUS_N_ Corelets, or **(E)** NPM1 Corelets. In each case, local activation drives local phase separation. (**F)** Concentration profiles across hnRNPA1 Corelets expressed in U2OS cell before and immediately after 2 min of local activation, showing local enhancement in 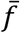, and depletion in the non-activated zone. **(G)** Simulations of locally activated spot of IDR-binding Core particles shows strong IDR enrichment, as observed in experiment. **(H)** Patterned activation examples with FUS_N_ Corelets. **(I)** Global activation causes droplets to condense in both cell nuclei. However, after local activation of two spots within the bottom cell, global activation of the entire cell does not initiate new nucleation events in that cell. Bars are 5 µm.

We also see this effect – phase separation activated with local, but not with global illumination-in Corelets formed from hnRNPA1_C_ (Figure 5C), FUS_N_ with extremely low 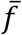 (Figure 5D), and the nucleolar protein NPM1 (Figure 5E). Since smaller activation zones are associated with a higher valence, this effect becomes particularly strong for highly localized activation. Indeed, by focusing light on a single diffraction limited spot, we find that we can drive highly localized droplet condensation (Figure 5F); simulations with localized activation support the physical picture of diffusive capture and concentration amplification (Figure 5G). Using patterned activation light, individual droplets could be written into different locations in the nucleus, to form 3×3 matrices and other arbitrary shapes (Figure 5H; Movies S5A-B). Interestingly, FRAP experiments suggest that these localized droplets can exhibit significantly different internal molecular dynamics than those that form under global activation (Figure S6). Moreover, in some cases where we locally activate a small number of single droplets, subsequent uniform illumination does not result in additional droplet condensation throughout the nucleoplasm (Figure 5I). This is consistent with the decreased 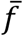 in these regions, which arises from activated Cores locally capturing IDRs and thereby depleting them from the non-activated regions; thus, patterned phase separation can impart “memory” into distal cytoplasm regions.

## Discussion

Our results provide the first true mapping of intracellular phase diagrams, which reveal a number of classical signatures associated with equilibrium phase diagrams, most remarkably droplet growth modes of nucleation and growth vs. spinodal decomposition, defined by the nested binodal and spinodal phase boundaries. These findings thus provide the first conclusive evidence that the concepts of equilibrium liquid-liquid phase separation are indeed applicable within living cells. And yet, living cells are certainly out-of-equilibrium systems, and our ability to map these intracellular phase diagrams raises many questions about the role of non-equilibrium activity in liquid-liquid phase separation. Indeed, while we use the Corelet system to show that the N-terminal IDR of FUS (FUS_N_) exhibits clear signatures of an equilibrium phase transition, this region of FUS is known to be subject to a number of post-translational modifications (PTMs) (Monahan et al. 2017). The FUS_N_ phase diagram thus must reflect a steady state interaction energy landscape, reflecting its average PTM state. These modifications can be dynamically modulated by cells in space and time, for example during development or through the cell cycle, providing the cell with a set of handles to dynamically structure these phase diagrams for particular functional requirements.

We have focused on using the Corelet system to examine the non-equilibrium biophysics of patterned intracellular phase transitions, revealing a powerful mechanism by which slowly diffusing multivalent complexes can capture and amplify the concentration of associated IDR binding partners, and thus drive local condensation. The ability to locally concentrate IDRs is particularly interesting, given that the phase diagram (Figure 3E) shows that even a modest degree of oligomerization, i.e. the binding of ∼4 IDRs in the case of FUS_N_, can promote phase separation. Thus, locally activating multivalent interactions, for example through protein phosphorylation by spatially-patterned kinases or the transcription of RNA, may be sufficient to drive local droplet condensation, even under conditions where the ligand (e.g. IDPs) are too dilute to phase separate upon global activation (Figure 6).

**Figure 6.**
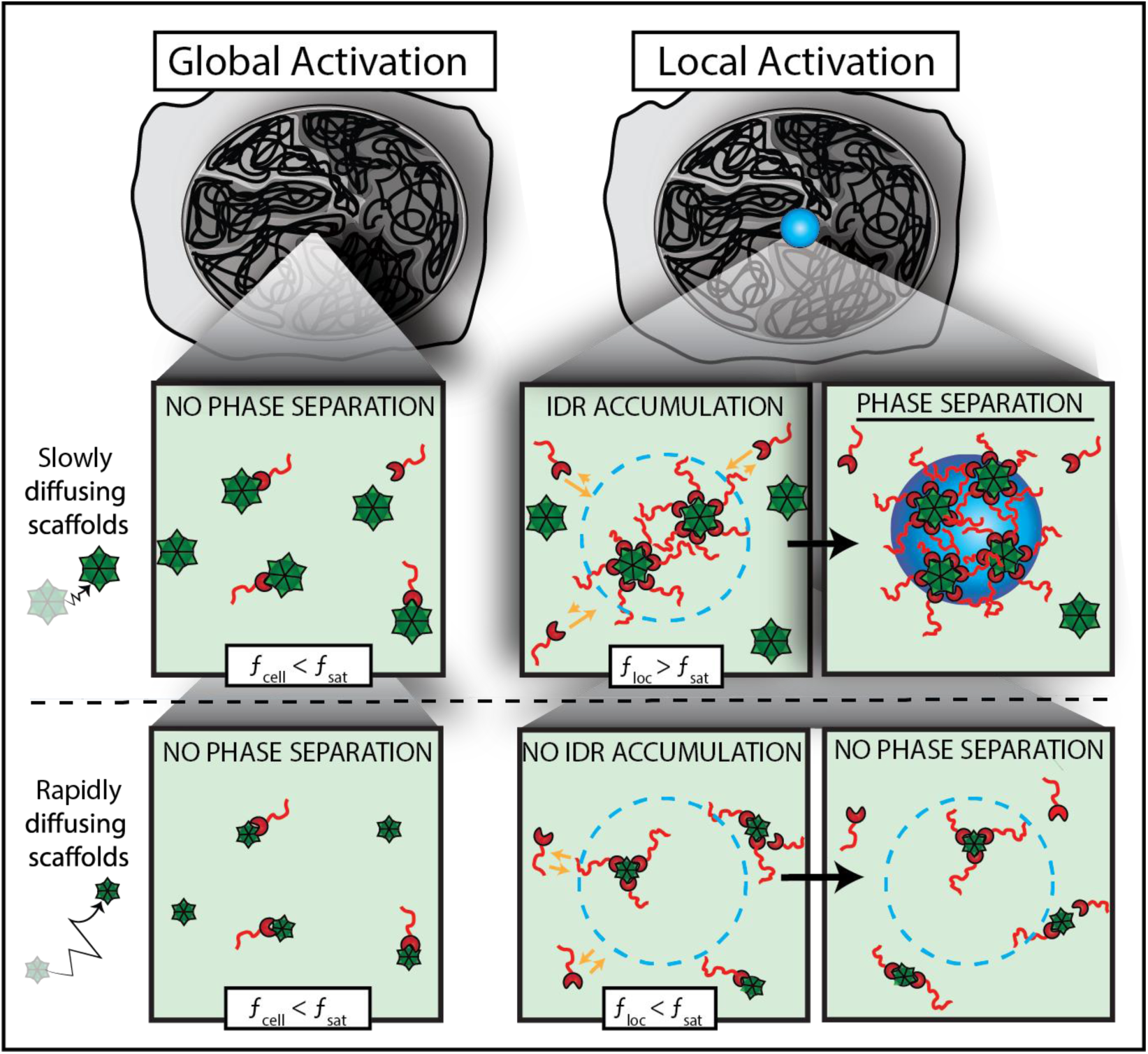
Model for localized phase-separation via diffusion capture. Schematic illustration showing how local activation can drive a diffusive flux of IDR towards slowly diffusing cores/scaffolds at the activation zone, causing high local valency, *f*_*loc*_, that exceeds the saturation threshold, *f*_*sat*_, for phase separation. Rapidly diffusing cores/scaffolds however, quickly diffuse away from the activated region and therefore *f*_*loc*_ does not cross *f*_*sat*_ and phase separation does not take place.

This diffusive capture mechanism may be particularly relevant for phase transitions involving nucleic acids, which are key components of many native intracellular condensates. DNA, mRNA and lncRNA often exhibit extremely slow diffusion rates <1µm_2_/sec, and together with their ability to simultaneously bind multiple disordered proteins, would allow them to serve as potent nucleators of local phase separation. Diffusive capture is thus likely to be a central mechanism in the emerging concept that liquid-liquid phase separation is involved in chromatin compaction and transcriptional control (Larson et al. 2017; Strom et al. 2017; Hnisz et al. 2017; Kwon et al. 2014; Berry et al. 2015), which involve a dynamic collection of numerous types of DNA and RNA. Indeed, while associated IDRs and other condensation-promoting ligands are often not present at particularly high concentrations (Biggin 2011), they are nevertheless known to bind to multivalent nucleic acids, which are distributed at relatively high concentrations throughout the nucleus.

The biomimetic Corelet system that we have used to elucidate patterned phase separation will provide a powerful tool for elucidating other aspects of the physics of condensed intracellular phases, and will serve to inspire other optogenetic nucleation platforms utilizing different multivalent core particles, or linear variants. These tools will find a broad range of uses, not only for interrogating fundamental cell biological questions, but also for synthetic biomaterials and organelle engineering applications. These approaches will increasingly synergize with those in materials science, for example in the design of bio-interfacing materials with novel properties arising from star-polymer architectures (Ren et al. 2016). Bioengineering of such structures and their interplay with fundamental studies on the non-equilibrium biophysics of intracellular phase transitions promises to be a fruitful area of future research.

We thank Yongdae Shin, Gena Whitney, Eje Chang, Carlos Chen, David Sanders and other members of the Brangwynne laboratory for help with experiments and comments on the manuscript. We also thank Michael Elbaum, Sam Safran and Rohit Pappu for helpful discussions. This work was supported by an HHMI Faculty Scholar Award, and grants from the NIH 4D Nucleome Program (U01 DA040601), DARPA (HR0011-17-2-0010), the Princeton Center for Complex Materials, an NSF supported MRSEC (DMR 1420541), as well an NSF CAREER award (1253035), and the US-Israel Binational Science foundation (2016508). D.B. acknowledges support through a Cross-Disciplinary Postdoctoral Fellowship from the Human Frontiers Science Program.

