## Supplementary Materials for "Mapping local and global liquid-liquid phase behavior in living cells using light-activated multivalent seeds"

#### Materials and methods

##### Cloning

DNA constructs used for tissue culture (Table S1) were cloned using In-Fusion HD cloning kit (Clonetech) in a standard reaction mixture comprising 20 ng of each 1-3 PCR-amplified inserts and 40 ng linearized pHR-SFFV backbone in a 5 µl reaction set to $50^{0}$C for 15 min. PCR fragments were produced using a standard PCR reaction using Phusion® High-Fidelity DNA Polymerase (NEB). Oligonucleotides were synthesized by IDT. PCR templates are listed in table S2. NLS sequence from Gallus gallus ferritoid (Bellapadrona and Elbaum, 2014) was incorporated by sequential PCR reactions. PCR products were purified using PCR purification kit (Qiagen) and verified on an agarose gel. pHR-SFFV backbone was linearized using HF-BamHI and HF-NOTI (NEB) following the manufacturer's instructions. Plasmids were transformed into Stellar cells (Clontech), from which single colonies were picked, grown in LB supplemented by Ampicillin overnight, and minipreped (Qiagen) following manufacturer instructions. All cloning products were confirmed by sequencing (GENEWIZ).

| # | Construct | Vector | promoter | Cell/organism |
|---|---|---|---|---|
| 1 | NLS-iLID-EGFP-FTH1 | pHR | SFFV | NIH3T3, U2OS, HEK293, HeLa |
| 2 | iLID-EGFP-FTH1 | pHR | SFFV | HEK293 |
| 3 | iLID-EGFP-FTH1 | pRS-2µ | TEF1 | *S. cerevisiae* |
| 4 | iLID-mtagBFP2-FTH1 | pcfj-150 | dao5 | *C. elegans* |
| 5 | FUS_N-mCherry-SspB | pHR | SFFV | NIH3T3, U2OS, HEK293 |
| 6 | FUS-mCherry-SspB | pHR | SFFV | U2OS |
| 7 | hnRNPA1_C-mCherry-SspB | pHR | SFFV | U2OS |
| 8 | DDX4_N-mCherry-SspB | pHR | SFFV | HEK293 |
| 9 | TDP43_C-mCherry-SspB | pHR | SFFV | HEK293 |
| 10 | NPM1-mtagBFP2 | pHR | SFFV | HeLa |
| 11 | FUS_N-mtagBFP2 | pHR | SFFV | U2OS |
| 12 | mCherry-SspB | pHR | SFFV | NIH3T3 |
| 13 | GFP-P2A-mCherry | pHR | SFFV | HEK293 |
| 14 | FUS_N-mCherry-SspB | pRS-2µ | TDH3 | *S. cerevisiae* |
| 15 | PGL-1-mCherry-SspB | pDEST-II R4 R3 | Endogenous | *C. elegans* |
| 16 | PGL-1-GFP | pDEST-II R4 R3 | Endogenous | *C. elegans* |

**Table S1 - List of constructs used in this study**

**Construction of stable and transient cell lines**

Corelet construct-containing Lentiviruses were produced by cotransfecting HEK293T cells plated on a 6-well plate for 24 hrs with the desired DNA constructs (1.25 µg), pCMVdR8.91 (1.1 µg), and pMD2.G (0.15 µg) using Lipofectamine™ 3000 (Invitrogen) following manufacturer instructions. 2 mL of viral supernatants were collected and filtered from cell debris using 0.45 µm filter (Fisher Scientific) within 2-3 days following transfection. HEK293, U2OS, NIH3T3, and HeLa cells were transduced while at 60% confluency on 6-well plates by adding 0.3-1.5 mL of the harvested Lentiviruses to the cell medium. Cell were cultured in 10% FBS (Atlanta Biological) DMEM (GIBCO) supplemented with penicillin and streptomycin at $37^0$C with 5% $CO_2$ in a humidified incubator. For transient expression, vectors were transfected into NIH3T3 cells as described above and imaged during the following 2-6 days. 1 day prior to imaging, cultured cells were trypsinized, quenched with medium, and plated on a 35-mm glass-bottom dish (MatTek) pre-coated for 20 min with 0.25 mg/ml fibronectin (Thermo).

**Construction and imaging of Corelet-expressing *S. cerevisiae***

NLS-free Core and $FUS_N$ IDR constructs were cloned into pRS-2µ vectors under TEF1 and TDH3 promoters, ACT1 and ADH1 terminators, and with LEU2 and URA3 markers, respectively. Both plasmids were simultaneously transformed into the CEN.PK2 strain of *Saccharomyces cerevisiae* using standard lithium acetate protocols.

For live-yeast imaging, single colonies were selected and grown in synthetic complete medium with 2% (w/v) glucose without leucine or uracil supplements (SC-Leu-Ura) overnight at 30 °C, reseeded in SC-Leu-Ura media at $OD_{600}$ of approximately 1.0 and grown at 30 °C until mid-log phase ($OD_{600}$ = 2.5-3.0) for imaging. Cells were briefly spun down at 3000 RPM for 5 min. The pellet was lightly resuspended in the supernatant; 4 µl was applied to a glass objective and sealed with a cover slip.

| # | Protein | Template | Fragment |
|---|---|---|---|
| 1 | iLID | AddGene #60413 | Full |
| 2 | SspB | AddGene #60415 | Full |
| 3 | FTH1 (Human) | HEK293 CDNA library | Full |
| 4 | $FUS_N$ (Human) | Shin et al., 2017 | 1-214 |
| 5 | $hnRNPA1_C$ (Human) | Shin et al., 2017 | 186-320 |
| 6 | $DDX4_N$ (Human) | Shin et al., 2017 | 1-236 |
| 7 | $TDP43_C$ (Human) | HEK293 CDNA library | 218-414 |
| 8 | NPM1 (Human) | Human NPM1 Gene cDNA clone plasmid (Sino Biological) | Full |
| 9 | GFP | Shin et al., 2017 | Full |
| 10 | mCherry | Shin et al., 2017 | Full |
| 11 | mtagBFP2 | Synthesized (IDT) | Full |
| 12 | PGL-1 (C. elegans) | Genomic DNA | Full |

**Table S2 – DNA templates and protein fragments used in this study.**

**Construction of Corelet expressing *C.elegans***

*C.elegans* were cultured according to standard methods. Worms expressing LOV2::mtagBFP2::FTH1 driven by dao5 promoter were established by mosSCI (Strain name CPB205, allele ptnIs136[dao-5p::LOV2::mtagBFP2::FTH1::tbb-2 3'UTR]. Worms expressing

PGL-1::mCherry::SspB (Strain name CPB207, allele ptnIs138[pgl-1p::pgl-1::mCherry::SspB]) and PGL-1::GFP (Strain name CPB 157, allele ptnIs92[pgl-1p::pgl-1::gfp(A206K)] were generated using CRISPR/Cas9 and driven by the endogenous promoter. CPB 205 and CPB 207 were crossed to create CPB 211[ptnIs136; ptnIs138], the PGL-1 Corelet line. CPB 211 was then crossed with CPB 157 to create CPB 212[ptnIs136; ptnIs138; ptnIs92], the line used to observe PGL-1::GFP recruitment to PGL-1 Corelets.

Worms were grown under standard conditions on OP50 bacterial lawns at 20 ºC. For imaging, embryos were dissected from gravid mothers in standard M9 buffer solution (3 g KH2PO4, 6 g Na2HPO4, 0.5 g NaCl, 1 g NH4Cl per 1 L) at room temperature, mounted on 3% agarose pads, and sealed with a cover slip.

**Live cell imaging**

Imaging was performed using an oil immersion objective (Plan Apo 60X/1.4, Nikon) on a laser scanning confocal microscope (Nikon A1) equipped with $CO_2$ microscope stage incubator at 5% $CO_2$ and 37$^0$C. Since EGFP and mtagBFP2 excitation overlaps with iLID activation spectrum, we performed pre-activation imaging as well as deactivation imaging (i.e. condensate disassembly) through mCherry channel only (560 nm excitation), which allowed visualization of only the IDR components. For global activation, cells were imaged sequentially by both mCherry and GFP (488 nm) channels, such that both Core and IDR components were visualized. Most activation protocols were conducted with excitation power of 0.1 nW/$\mu m^2$ measured with an optical power meter (PM100D, Thorlabs). mTagBFP2 labeled constructs were imaged similarly through 405 nm excitation channel. Global activation protocols were performed with dual channel imaging (568 nm and 488 nm) with frame intervals of 4.4 s and 2.2 s for 120x120 $\mu m^2$ (1024x1024 pix) and 60x60 $\mu m^2$ (512x512 pix) frame sizes respectively. In the case where a fast frame rate was desirable (Figure 3G-I), a frame interval of 0.5 s was used (264x264 pix). Local activation was performed by activating a pre-defined ROI through stimulation mode at either 488 nm or 405 nm wavelength. FRAP experiments were performed similarly to local activation, yet at an ROI size dictated by diffraction limit and higher activation power (~100 fold).

**Estimation of endogenous and exogenous ferritin cellular levels**

Untransfected HEK293 cells and those stably expressing $FUS_N$ Corelets were grown to approximately 90% confluency in 60 mm plates. Cells were trypsinized and collected in PBS with protease inhibitor. Cell lysates were prepared by sonification and protein concentration was quantified with Bradford Assay. Samples of 0.4 µg/µL whole cell lysate were prepared in Novex NuPAGE lithium dodecyl sulfate (LDS) buffer (Invitrogen) supplemented with 75 mM dithiothreitol (DTT; Thermo Scientific) as a reducing agent. FTH1 and FTL1 recombinant protein standards (ProSpec) were diluted to 100 ng in 25 µl of the same LDS/DTT buffer. After briefly boiling all samples at 100 °C, 25 µl (10 µg protein weight cell lysate, 100 ng protein standard) of denatured sample was loaded to NuPAGE 4-12% Bis-Tris protein gel and run with NuPAGE MOPS buffer (Invitrogen) at 100 V for 90 minutes. Wet transfer to a Polyvinylidene difluoride membrane was performed at 30 V for 1 hour in NuPage Transfer buffer (Invitrogen). The membrane was blocked with 5% Non-Fat Dry Milk (Nestle) in TBST. The β-Actin strip was cut from the membrane directly above the 30 kDa and 50 kDa protein standards and probed with rabbit anti-β-Actin (ab8227, abcam) overnight at 4 °C. The remaining membrane was probed under the same conditions with mouse anti-FTH1 (MABC602, Millipore Sigma). After washing with TBST, the β-Actin strip was probed with anti-rabbit horseradish peroxidase (HRP) (111-035-144, Jackson ImmunoResearch), and the FTH1-probed membranes were probed with anti-mouse HRP (115-035-062, Jackson ImmunoResearch), both at room temperature for 30 minutes. Chemiluminescence was induced with SuperSignal West Pico substrate (Thermo Scientific) and imaged with a ChemiDoc MP Imaging System (BioRad).

**Determining diffusion coefficients and absolute concentrations for Corelets components**
Data for diffusion and concentration of proteins were obtained using fluorescence correlation spectroscopy (FCS). The measurements were performed on U2OS cells expressing either GFP labeled cores or mCherry labeled IDR components using a laser scanning confocal microscope (Nikon A1) with an oil immersion objective (Plan Apo 60X/1.4, Nikon). All measurements and data analysis were performed using the SymPhoTime Software (PicoQuant).

The autocorrelation function for simple diffusion is:

$$G(\tau) = G(0)\left(1+\left(\frac{\tau}{\tau_D}\right)\right)^{-1}\left(1+\left(\frac{\tau}{\kappa^2\tau_D}\right)\right)^{-0.5}$$

Here, $G(0)$ is magnitude at short time scales, $\tau$ is the lag time, $\tau_D$ is the half decay time, and $\kappa$ is the ratio of axial to radial of measurement volume. The parameters $\tau_D$ and $G(0)$ are optimized in the fit and are used to determine the diffusion coefficient and molecule concentration.

While mCherry fluorescence was converted to absolute concentration using the FCS measurement (Figure S7), GFP fluorescence conversion was done by determining the exact mCherry-to-GFP fluorescence ratio while using the mCherry fluorescence to concentration ratio as a set point. mCherry-to-GFP fluorescence ratio was determined by equi-molar expression of mCherry and GFP monomers in HEK293 cell using the auto-catalytic P2A containing construct mCherry-P2A-EGFP, which unlinked the two proteins such that FRET is prevented.

**Image analysis and phase diagram construction**
Nucleoplasm boundaries not including nucleolar regions were determined based on the Core component partitioning (EGFP channel) by applying an automated image segmentation Matlab code. Histograms of fluorescent signal were plotted as shown in Figure 3A-C. For each cell, the dilute phase concentration (Figure 3E, triangle) was determined according to the center of the left peak. This was done by calculating the median of fluorescent values having histogram amplitudes greater than half of the peak's maximal value. Dense phase concentration (Figure 3E, diamonds) was determined automatically by performing subsequent image segmentation to identify intra-nuclear condensed regions following by morphological etching of 1.1μm and averaging fluorescence readout in the remaining condensates area. Dense phase concentration in condensates smaller than 2.2 μm across have not been taken into consideration for phase diagram reconstruction. This threshold in condensates diameter was determined based on the observation that when plotting fluorescent profile across condensates, only condensates larger than this critical value displayed a clear plateau at their center (Figure 3A-C insets), presumably due to point spread function of the z-axis. The maximal amplitude of the right peak of the histograms (Figure 3A-C histograms), less reliably represent the mean dense phase concentration as it gives large weight to pixels located at droplet's periphery as well as to out of focus droplets. Nevertheless, the dense

phase values determined by fluorescent profiles were very similar to the center of the right peak. Mean nuclear concentrations of cores (Figure 3E, circles and asterisks) and IDR components were calculated by taking the mean fluorescence at the entire segmented nucleoplasm after 10 min of activation. While mean nucleoplasmic core concentration is fixed, nucleoplasmic levels of $FUS_N$-based IDR component as well as the mean valence, which is derived from it, continuously increase during activation. This occurs since $FUS_N$-based IDR component (but not $hnRNPA1_C$) have both cytoplasmic and nucleoplasmic subpopulation, where once activation is applied, the fast uptake and sharp depletion of nucleoplasmic IDR component monomers, drives net flow of cytoplasmic IDRs into the nucleus. For that reason, valence values were determined only after steady state is reached (t ~ 5 min). Contrary to the steady state binodal line, the spinodal line (i.e. the mechanism by which phase separation occurs) was determined accordingly to the valence at t = 0 (see Figure S5).

**Simulations**

We developed a simple Monte Carlo simulation of the Corelet system, using Matlab. Simulations were run with 500 "Core" particles, and 1200 "IDR" particles, randomly distributed in a 2D simulation space with reflecting boundary conditions. Particle diffusivity is modeled by introducing a random "kick" at each time step, Δt, such that particles move by an amount (in both x and y) given by $\sqrt{4D_i\Delta t}\cdot\xi$, where ξ is a normally distributed random variable with mean 0 and standard deviation 1. The particle diffusivity, $D_i$, was varied for different simulations with most simulations set at the experimentally-determined diffusivities, i.e. $D_{Core}$=3μm$^2$/sec, $D_{IDR}$=43.5μm$^2$/sec. At each time point, the position of each IDR particle is checked to see if it is within a set interaction distance of any Core particle. For particles within a defined activation zone, if the IDR is close enough to bind the Core, and the Core is not already saturated with a defined number of IDRs ("maxidrs"), then the IDR particles bind, by remaining at this fixed position relative to the Core particle position, while the Core particle (and thus associated bound IDR particles) continues with its diffusive motion, updated at each time step. For all simulations, if the diffusive motion of the Core particle takes it outside of the activation zone, then any bound IDR particles are released.

### Supplementary Figures

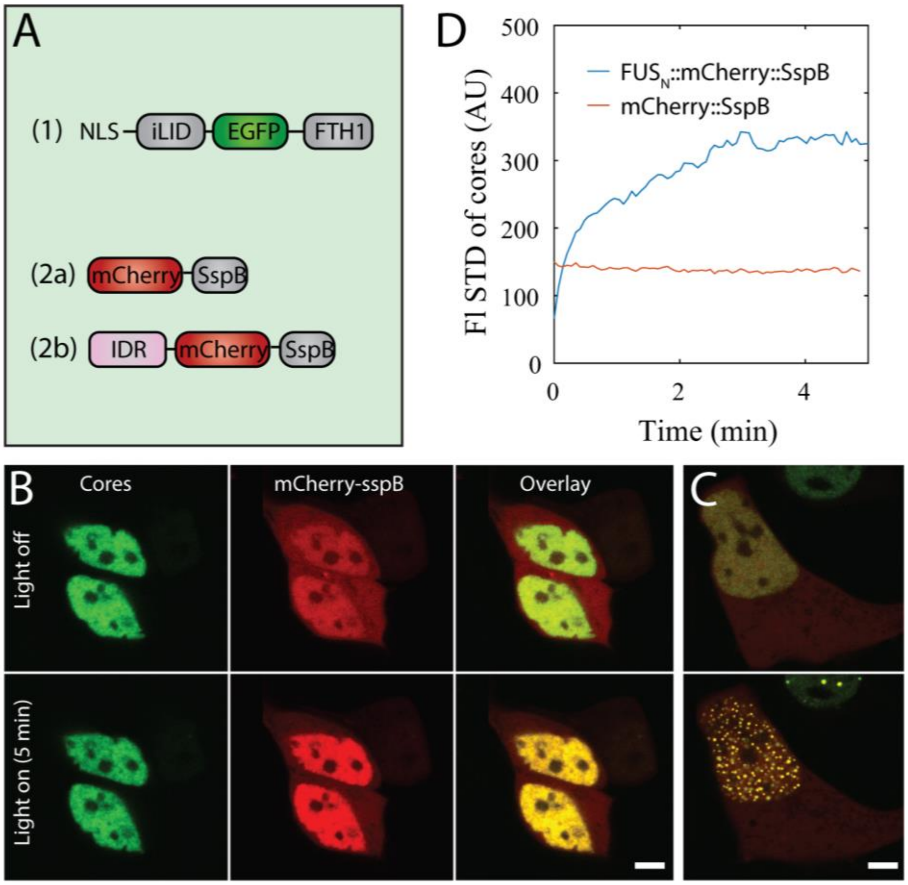


**Figure S1. IDR-free Corelets do not phase separate. (A)** Schematic diagram of Corelets with and without IDR fused to the SspB constructs. **(B)** Representative NIH3T3 cells transiently expressing core (green) and mCherry-SspB constructs (red) before and after 5 min photo-activation at 0.3 nW/μm$^2$, showing no puncta formation. The concentration of cores and SspB-to-core ratio (i.e. mean expected valence $f^{-1}$) are 18 μM and 0.4 respectively, which position the cell well within the $FUS_N$ Corelets bimodal region, where demixing occurs (see Figure 3D). Activation-driven enhancement in nuclear mCherry:SspB (red) is a signature of a sharp drop in nuclear levels of SspB monomers (i.e. unbound by cores), which is compensated by a net flux from the cytoplasmic reservoir. Activation-driven nuclear accumulation, which was observed in partitioning constructs such as $FUS_N$ and $DDX4_N$ but not in non-partitioning constructs such as FUSwt, $TDP43_C$, and $hnRNPA1_N$, indicate that SspB constructs are indeed capturing by cores, yet in the case of IDR-free SspB construct decoration of cores is insufficient for driving phase separation. In few cases we have observed 1-2 small dim spots in response to photo-activation, presumably as a result of weak mCherry or SspB homotypic dimerization. **(C)** $FUS_N$ Corelets with similar SspB-to-core ratio as in B undergoing phase separation under similar excitation conditions. **(D)** Standard deviation of the intensity of cores for SspB construct with or without $FUS_N$ (blue and red lines respectively) showing no change during activation time for IDR-free Corelets. Scale bars are 5 μm.

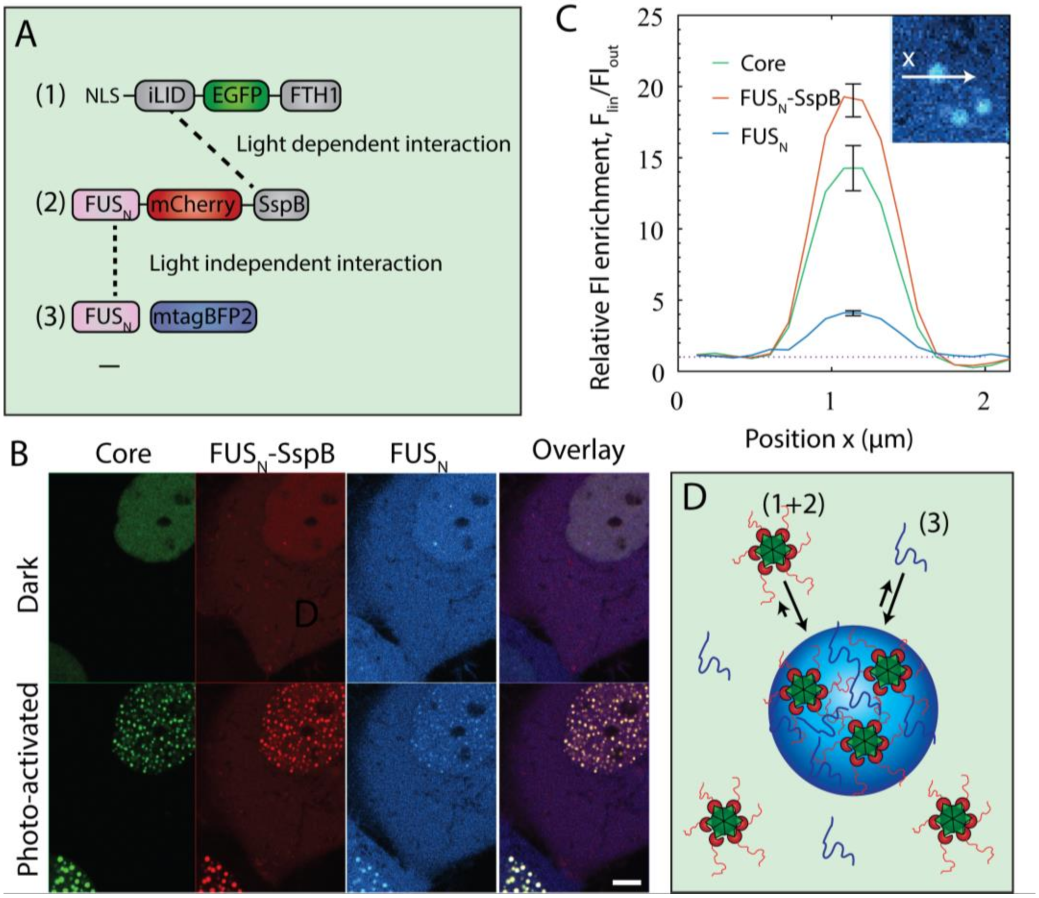


**Figure S2. Recruitment of SspB-free FUS$_N$ monomers by FUS$_N$ Corelets. (A)** Schematic diagrams of the two-component Corelet system as well as an SspB-free light-insensitive FUS$_N$ monomer. **(B)** Fluorescent images of stable HEK293 cells expressing Cores (green), FUS$_N$-SspB (red), and FUS$_N$ (blue) before and after 5 min of blue light activation, showing co localization of all three components. **(C)** Quantification of the relative enrichment of the three components inside droplets after 5 min of activation. **(D)** Schematic illustration showing recruitment of FUS$_N$ monomers through IDR-IDR interactions. Scale bar is 5 μm.

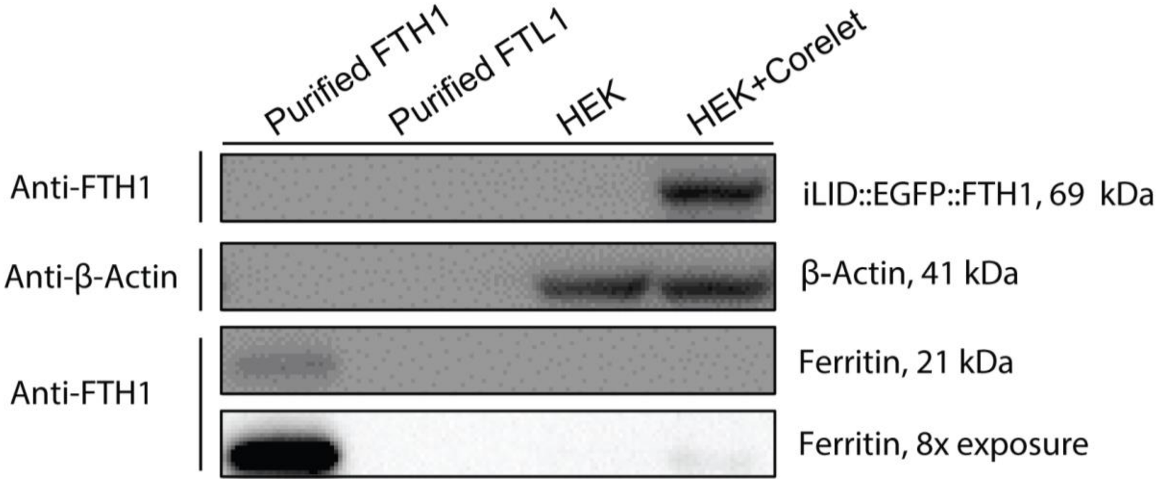


**Figure S3. Native ferritin does not interfere with Corelet formation**.
Western blot analysis of ferritin in Corelet expressing HEK293 cells (HEK+Corelets) show small amount of native FTH1 as compared to exogenous Core components. Untransfected cells (HEK) act as a negative control. 100 ng of purified heavy chain (FTH1) and light chain (FTL1) ferritin (ProSpec) loaded as positive and negative controls, respectively. Upper three panels imaged were imaged with the same exposure time (~10 sec) while the bottom panel is a replication of that directly above it but imaged with eight times longer exposure. β-Actin shown as an internal loading standard. Calculated molecular weight indicated and band identification is shown to the right, and antibody specificity used for each blot is indicated on the left.

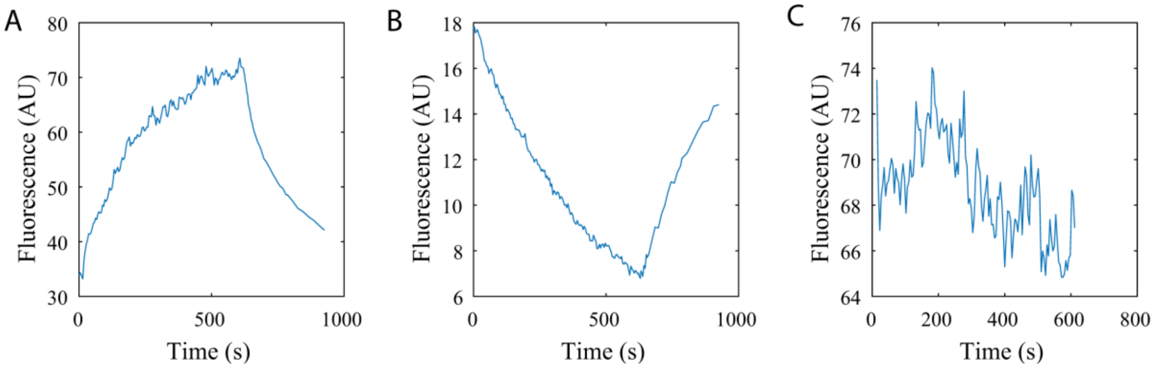


**Figure S4. Transitions in local concentration of IDR and Core components in $FUS_N$ Corelet expressing U2OS cells during activation. (A)** $FUS_N$ IDR concentration in nucleoplasm and **(A)** cytoplasm during 10 min activation and 5 min deactivation. Immediately after activation, the concentration in nucleoplasm begins increasing while the cytoplasmic concentration begins decreasing, suggesting that nucleoplasmic IDR components are quickly captured by cores, leading to a sharp drop in unbound IDRs in nucleoplasm, and a net nuclear influx of unbound IDRs from the cytoplasm. **(C)** Core component concentration in the nucleoplasm does not change significantly during 10 min activation and 5 min deactivation. We also do not observe any sharp drop during the first few seconds of activation, suggesting minimal FRET response between the EGFP of the Core component and the mCherry of the IDR component.

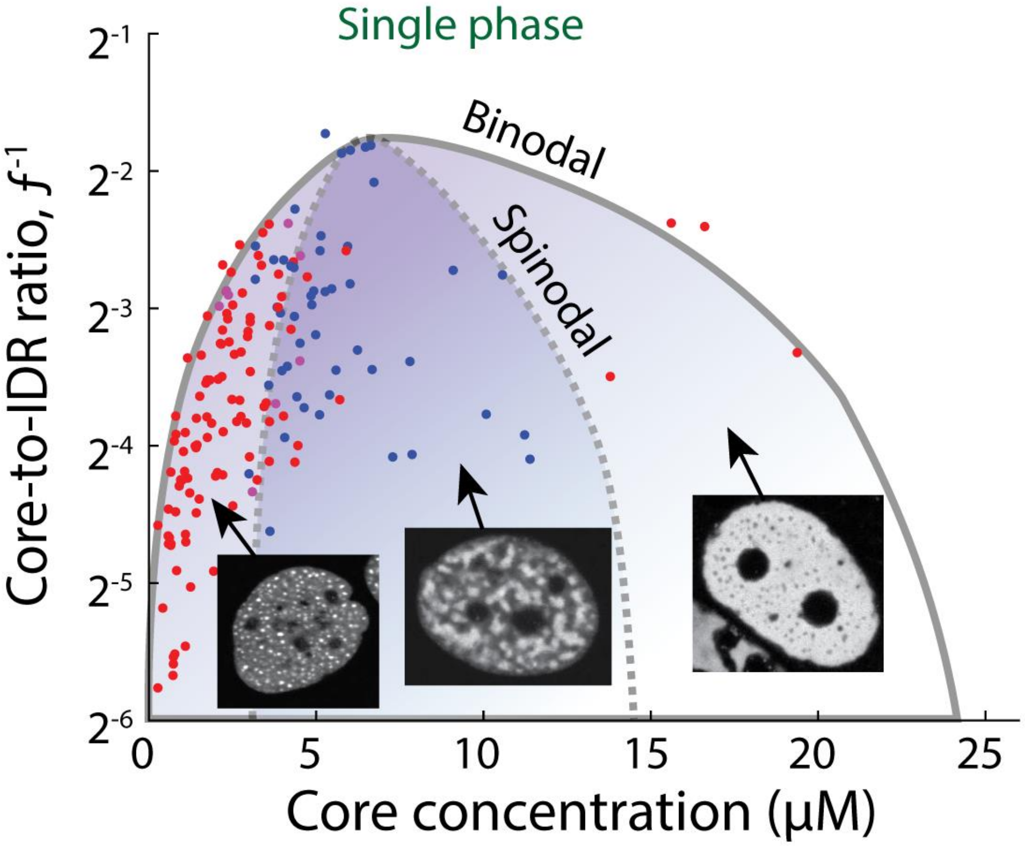


**Figure S5. Modes of $FUS_N$ Corelets condensation.** Phase diagram of $FUS_N$ Corelets, where red symbols indicate average nuclear concentrations for which phase separation follows nucleation and growth mechanism of dense phases (left) or dilute phases (right), and blue symbols are concentrations where spinodal decomposition is observed. Points represent mean fluorescence upon activation. Condensation mode was determined accordingly to the spatial fluorescence patterns appearing on the first imaging frame taken 2 sec after light activation. Purple symbols represent cells in which the exact condensation mode could not be conclusively determined. Dashed line represents estimated location of spinodal boundary. Note vertical axis on a log 2 scale.

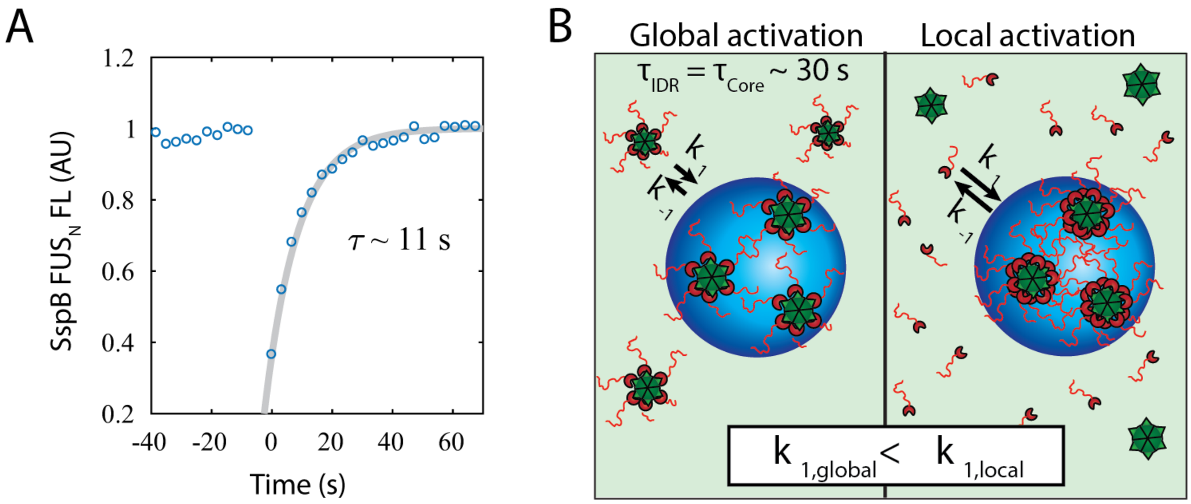


**Figure S6. Locally activated droplets are more dynamic than Globally activated droplets. (A)** Fluorescence recovery of mCherry labeled $FUS_N$ Corlets after photobleaching a 5 min long locally activated droplet. Droplet size reached steady state volume prior to FRAP. Measured recovery rate is roughly 3 times faster than in a globally activated cell (main text Figure 1D). **(B)** Schematic illustration showing the different dynamics in globally and locally activated cells. Note that locally activated cells typically reach higher condensed phase valency.

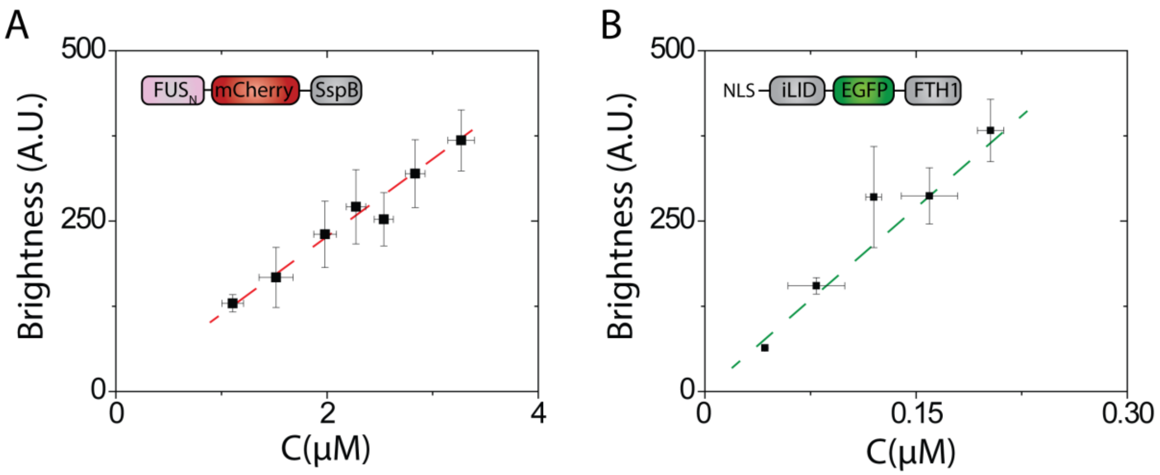


**Figure S7. Characterization of fluorescent protein quantum efficiency and microscope system detector sensitivity.** Plot of molecular brightness of **(A)** $FUS_N$-mch-SspB **(bB)** NLS-iLID-GFP-FTH1 obtained from laser scanning confocal microscope as a function of the protein concentration *C* measured in cellular nuclear plasma using fluorescence correlation spectroscopy. Data are collected on a Nikon A1 laser scanning confocal microscope with Plan Apo 60X/1.4 oil immersion objective and PicoQuant hybrid photomultiplier detector.

### Supplementary movies

**Movie S1–** $FUS_N$ Corelet-expressing (stable) U2OS cells photo-activated for 10 min. Cells exhibiting either nucleation and growth or spinodal decomposition type dynamics. Red and Green channels show IDR and Core components respectively. Upon deactivation, dissolving condensates are monitored through IDR channel only.

**Movie S2A** – $FUS_N$ Corelets expressing U2OS cells (stable) undergoing nucleation and growth of dense phases upon 10 min activation.

**Movie S2B** – High frame rate confocal imaging of $FUS_N$ Corelets expressing U2OS cells (stable) undergoing spinodal decomposition of dense phases upon 2 min activation.

**Movie S2C**– High frame rate confocal imaging of $FUS_N$ Corelets expressing U2OS cells (stable) undergoing spinodal decomposition of dilute phases upon 2 min activation.

**Movie S2D** – High frame rate confocal imaging of $FUS_N$ Corelets expressing U2OS cells (stable) undergoing nucleation and growth of dilute phases upon 4 min activation.

**Movie S3A** – Simulation of half-cell activation at high power (24 available IDR binding sites per core) with $D_{IDR} = 43.5\ \mu m^2/s$ and $D_{Core} = 3\ \mu m^2/s$, similar to Corelet system, showing concentration build-up of IDRs at the interface between activated and non-activated zones (left and right sides, respectively).

**Movie S3B** – Simulation of half-cell activation at low power (1 IDR binding site per core) with $D_{IDR} = 43.5\ \mu m^2/s$ and $D_{Core} = 3\ \mu m^2/s$, similar to Corelet system, showing similar IDR concentration across the activated zone (left side).

**Movie S4** – Multiple activation-deactivation cycles applied on a decreasing fraction of a $FUS_N$ Corelets expressing U2OS (stable) cell.

**Movie S5A** – Applying activation pattern of a 3x3 array of single droplets on an NIH3T3 $FUS_N$ Corelets expressing (transient) cell.

**Movie S5B** – Applying an arbitrary activation pattern on an NIH3T3 $FUS_N$ Corelets expressing (transient) cell.
